# Evidence-based gene expression modulation correlates with transposable element knock-down

**DOI:** 10.1101/2020.08.15.252312

**Authors:** José Hernandes-Lopes, Danielle M. Quintanilha, Erika M. de Jesus, Fabrício M. Lopes, Raphael B. Parmigiani, Bruno Karolski, Henrique M. Dias, Thomas B. Jacobs, Anamaria A. Camargo, João P. Kitajima, Marie-Anne Van Sluys

**Affiliations:** Departamento de Botânica, Universidade de São Paulo, São Paulo, Brazil; Universidade Federal Tecnológica do Paraná, Cornélio Procópio, Brazil; Ludwig Institute for Cancer Research, São Paulo, Brazil; Idengene Medicina Diagnóstica, São Paulo, Brazil; Laboratório de Simulação e Controle de Processos (LSCP-POLI-USP), São Paulo, Brazil; Center for Plant Systems Biology, VIB, Ghent, Belgium; Centro de Oncologia Molecular, Hospital Sírio-Libanês, São Paulo, Brazil; Mendelics Análise Genômica, São Paulo, Brazil

**Keywords:** ethylene, transcriptome, retrotransposon, Tnt1, tobacco, RNAi

## Abstract

**Background:** Transposable elements (TEs) are major components of plant genomes. Despite being regarded as “junk DNA” at first, TEs play important roles for the organisms they are found in. The most obvious and easily recognizable effects caused by TEs result from their mobility, which can disrupt coding sequences or promoter regions. However, with the recent advances in transcriptomics, it is becoming increasingly evident that TEs can act as an additional layer of gene expression regulation through a number of processes, which can involve production of non-coding RNAs. Here, we describe how Tnt1, a stress-responsive LTR-retrotransposon, interferes with gene expression and modulate a number of developmental aspects in tobacco.

**Results:** Through an RNAi approach, we generated tobacco (HP) lines knocked-down for Tnt1 expression. Quantitative RT-PCR experiments confirm that Tnt1 is downregulated in HP lines after ethylene exposure. A RNA-seq experiment was performed and through two independent bioinformatic approaches (with different stringencies) we found 932 and 97 differentially expressed genes in HP lines. A number of phenotypes were observed in such lines, namely lesion mimicry in leaves, underdevelopment of the root system, overproduction of root hairs and early loss of seed viability. Folding prediction of part of the Tnt1 mRNA reveals putative stem-loop secondary structures containing transcriptional regulation sequences, suggesting it could be a source of small RNAs. We also propose a model to explain the Tnt1 expression in both homeostatic and stress conditions, and how it could interact with stress-responsive genes.

**Conclusions:** Our results are consistent that interferences with Tnt1 transcript levels correlate with transcriptomic and phenotypic changes, suggesting a functional role for this element during plant development and stress response.

## BACKGROUND

Most genomes harbor a particular type of genetic elements collectively known as Transposable Elements (TE). TEs encode proteins that enable their own mobilization within the genome. Discovered by Nobel laureate Barbara McClintock in the 1940’s, TEs were first described as “controlling elements” since mobilization of specific TEs (*Ac* and *Ds*) in maize caused a variegated phenotype in kernels due to an insertion into the C locus that is responsible for anthocyanin synthesis [1]. Despite this, TEs in eukaryotes were at first widely regarded as “junk DNA” or “selfish genes” due to their self-replicating nature, mutagenic potential and the lack of an obvious function for the host genome [2].

The idea of TEs as simply being genomic parasites was gradually abandoned with the ever-increasing understanding of eukaryotic genome structure. TEs are known to be found in virtually all domains of life [3]. Whereas TEs make up about 45% of the human genome [4,5], the TE-derived content can be much higher in domesticated plants such as sorghum (~62%), tomato (~63%), wheat (~80%) and maize (~85%) [6, 7]. Nowadays it is well accepted that TEs are able to generate genome structural and functional variation as a result of their mobile nature and predisposition to recruit epigenetic silencing mechanisms. Through a number of processes, they can deeply affect epigenetic variation, alter or create new gene regulatory networks, as well as the formation of new proteins through the spreading of functional motifs [8, 9, 10].

Given the potential deleterious effects of transposition, expression and mobility of TEs are usually tightly controlled in eukaryotes. Modifications of histone tails, DNA methylation and alterations in chromatin packing and condensation are amongst the most well-known mechanisms involved in TEs silencing, but there are examples of post-transcriptional silencing of TEs by RNA interference (RNAi) (see 11 for a review). Interestingly, in vertebrates silencing can be achieved by DNA editing through APOBEC enzymes, which selectively edit the promoter region of LTR type retroelements [12, 13]. Thus, while deleterious effects of new insertions are negatively selected, advantageous changes can be incorporated (for reviews, see reference 9 and 14), playing an important role in evolution. Such is the case of maize cultivars, which are living examples of genome evolution driven by transposable elements [15].

Besides the classically-accepted role as drivers of genetic diversity, stress-related expression of TEs is also demonstrated to participate in different regulatory pathways, such as the human *Alu* element, which seems to regulate protein translation after exposure to stress [16]. While there are many examples of stress-induced expression of TEs, recent studies also recognize TEs as important components for the maintenance of biological processes. Also, TEs have been shown to be a significant source of noncoding RNAs and to interfere in the small RNA (sRNA) machinery, which are key regulators of gene expression in plants [17, 18]. An example of such interference is the expression of the TE MIKKI during rice root development. MIKKI transcripts act as decoys for miR171, which usually targets and silences *OsSCL21.* By mimicking *OsSCL21*, MIKKI sequesters miR171 molecules, culminating in *OsSCL21* upregulation [19].

The Tnt1 retrotransposon is an interesting transcriptionally-active TE in somatic tissues of *Nicotiana tabacum* plants growing under normal conditions [20, 21]. Tnt1 (Genbank: X13777) is a multicopy Long Terminal Repeat (LTR) retrotransposon that was first discovered after its insertion in the nitrate reductase coding sequence [22]. It is estimated to have more than 600 insertions in the tobacco genome [23] and it also has homologs in other Solanaceae under different names (*e.g.* Retrolyc1/TLC1 in tomato and Retrosol in potato) [24–26].

Being an LTR element, Tnt1 is composed of central open-reading frames (ORFs) flanked by 5’ and 3’ LTRs, which can be further divided in U3, R and U5 regions [27]. In tobacco, Tnt1 insertions are classified in three subfamilies (Tnt1A, Tnt1B and Tnt1C), which have highly conserved sequences except for their U3 regions [28]. Regulatory sequences present in each of the U3 are responsive to different hormone induction [20, 21, 29]. Interestingly, the U3 region of Tnt1A (U3A) shows sequence repeats highly similar to the GCC core, present in the promoter region of ethylene-responsive genes [30, 31]. These results suggest that expression of this subfamily could also be induced by ethylene, but they remain to be experimentally validated. Expression of Tnt1 is also known to occur in homeostatic conditions. During normal development, expression of Tnt1 was reported in roots, leaves and petals [20]. Although interesting, more data on this basal expression is lacking in the literature, and whether it has a role for tobacco development or represent a residual escape from endogenous silencing mechanisms is still an open question.

Given the ever-increasing number of proposed mechanisms through which TEs can exert important functions, either at genome or organism levels, Tnt1 is an interesting target for functional studies. Thus, this work aims to understand the Tnt1 pattern of expression and its potential role during plant development. Through an RNAi approach we observed phenotypic changes caused by Tnt1 downregulation, such as increased root hair production, underdevelopment of the root system and decreased seed viability. Transcriptome profiling of downregulated Tnt1 plants revealed the dynamics of Tnt1 expression and a close association with ethylene biosynthesis and responsive genes in tobacco. Taken together, these results reveal the importance of the Tnt1 retroelement for normal tobacco development.

## RESULTS

### Tnt1 expression knockdown reveals a connection with ethylene biosynthesis and responsive genes

Given the presence of GCC-like motifs in the U3A region of Tnt1A insertions, we first tested if expression of Tnt1 is upregulated upon ethylene stimulus. In parallel, to understand if perturbations in the level of Tnt1 transcripts would have any detectable effect, we developed transgenic tobacco lines expressing an RNAi (hairpin, HP) construction under the Cauliflower mosaic virus 35S promoter, targeting the Tnt1 reverse transcriptase (RT) domain (Additional file: Figure S1). These lines are herein called HP(x), where (x) corresponds to an independent transformation event, followed by T(n), where (n) correspond to the generation of the transgenic line (T0 = the plant regenerated from the callus).

We treated two WT and two HP1 (T2) plants with ethylene, while two other individuals of each genotype did not receive the treatment (control group). After 24 hours of treatment we quantified the expression (quantitative RT-PCR) of three different Tnt1 coding domains in all individuals. Among individuals that were not exposed to ethylene treatment, HP1 plants had higher expression level of all three Tnt1 domains than the WT (Figure 1A). However, upon ethylene treatment both WT and HP1 plants overexpressed the three Tnt1 domains when compared to the control group, demonstrating that this gaseous hormone indeed induces Tnt1 expression. Interestingly, while WT plants increased Tnt1 expression by a factor of 35 to 50, HP1 plants had an increase of around 20-fold. These qRT-PCR results are consistent with decrease of Tnt1 expression in HP lines treated with ethylene, confirming the knock-down of Tnt1 by the RNAi construct. Since ethylene induced Tnt1 expression and it was upregulated in untreated HP1 plants, we asked whether HP plants produced more ethylene under normal conditions. Thus, we tested ethylene emissions in 90-day-old WT, HP1 (T1), HP1 (T2) and HP8 (T1) plants using gas chromatography. All HP samples showed a significant increase in ethylene production when compared to WT (Figure 1B).

**Figure 1.**
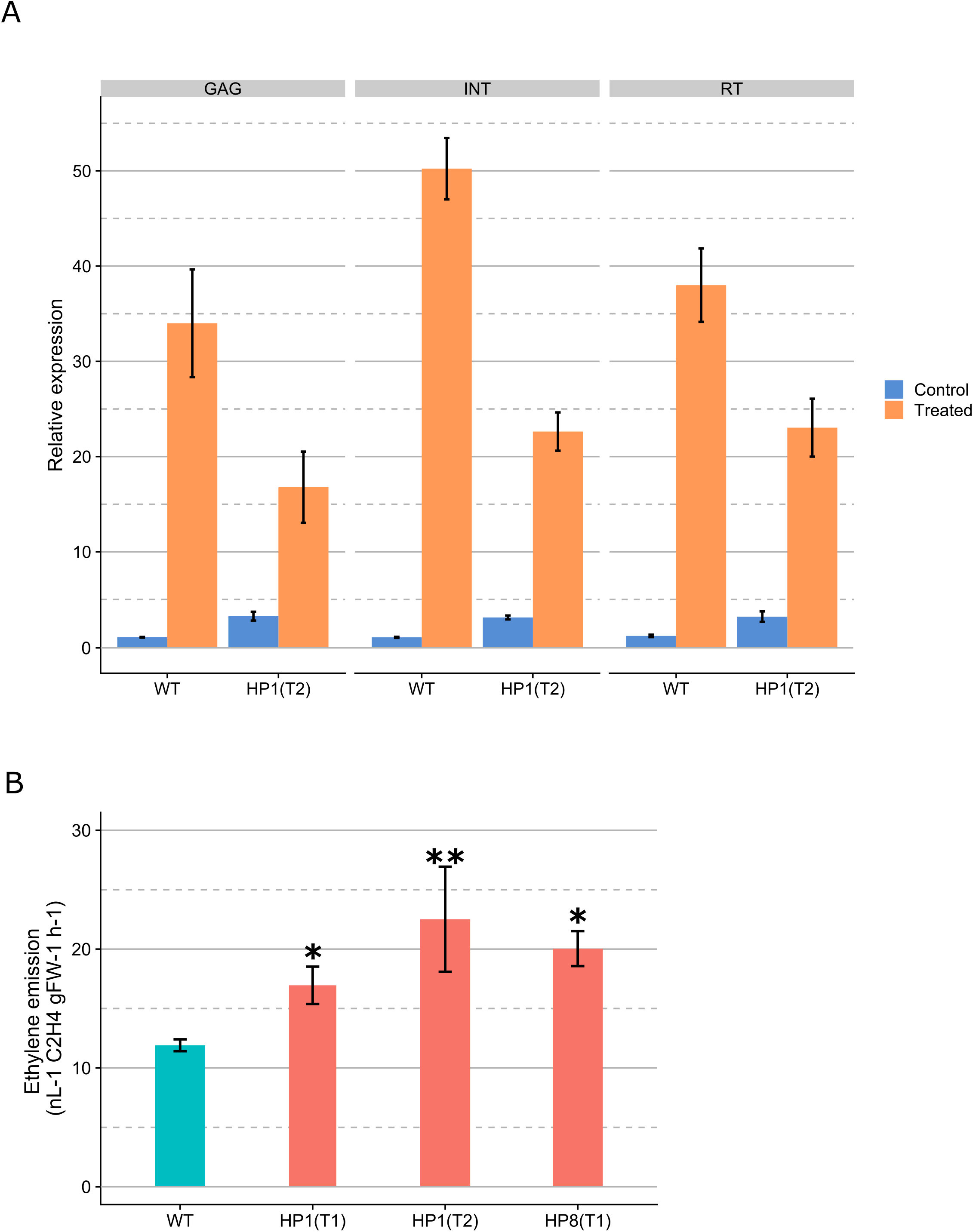
Dynamics of Tnt1 and ethylene emission in WT and HP lines. **(A)** Relative expression of three Tnt1 domains in 60-days-old plants measured by qRT-PCR. Tnt1 expression is induced upon exogenous ethylene application (10 μL/mL for 24hs). Control plants were incubated in sealed containers without input of ethylene. **(B)** Ethylene emission in 90-day-old HP plants measured by gas chromatography. Bars indicate standard error. Significance of the difference between HP and WT was assessed by a one-tailed Student’s t-test: (**) significant at the 0.004 level; (*) significant at the 0.017 level.

Because ethylene biosynthesis is known to follow a circadian cycle, we also investigated the expression dynamics of Tnt1 and some selected ethylene biosynthesis and responsive genes in WT plants throughout a period of 48 hours. For these, we used 15-day-old plants grown under a 12-hours-light / 12-hours-dark regime. Expression of two genes related to the circadian clock in *Nicotiana* species confirmed synchronization of the samples (*NtCP-23* and *NtTOC1*, reaching highest levels of expression at 12PM and 6PM respectively; Additional file 1: Figure S2A).

As expected, the ethylene biosynthetic genes *ACO1* and *ACO2* were transcribed in a circadian fashion in WT plants, peaking at the beginning of the light period (6h; Figure 2A). Accordingly, the ethylene responsive genes *ER24, JERF1* and *TEIL* also presented a circadian cycle: while *ER24* is consistently more expressed at noon, both *JERF1* and *TEIL* reached highest expression values at 6h, much like both *ACO1* and *ACO2* (Figure 2B). Expression of Tnt1 in the WT (evaluated by the RT domain) however, showed no signs of a circadian rhythm, presenting great variation in expression between biological replicates (Figure 2C). We then checked specific expression of the U3A region (expected to be responsive to ethylene). When biological replicates are plotted separately, it becomes clear that expression of Tnt1A is indeed decoupled from the circadian cycle as each sample showed different levels of U3A expression (Figure 2C).

**Figure 2.**
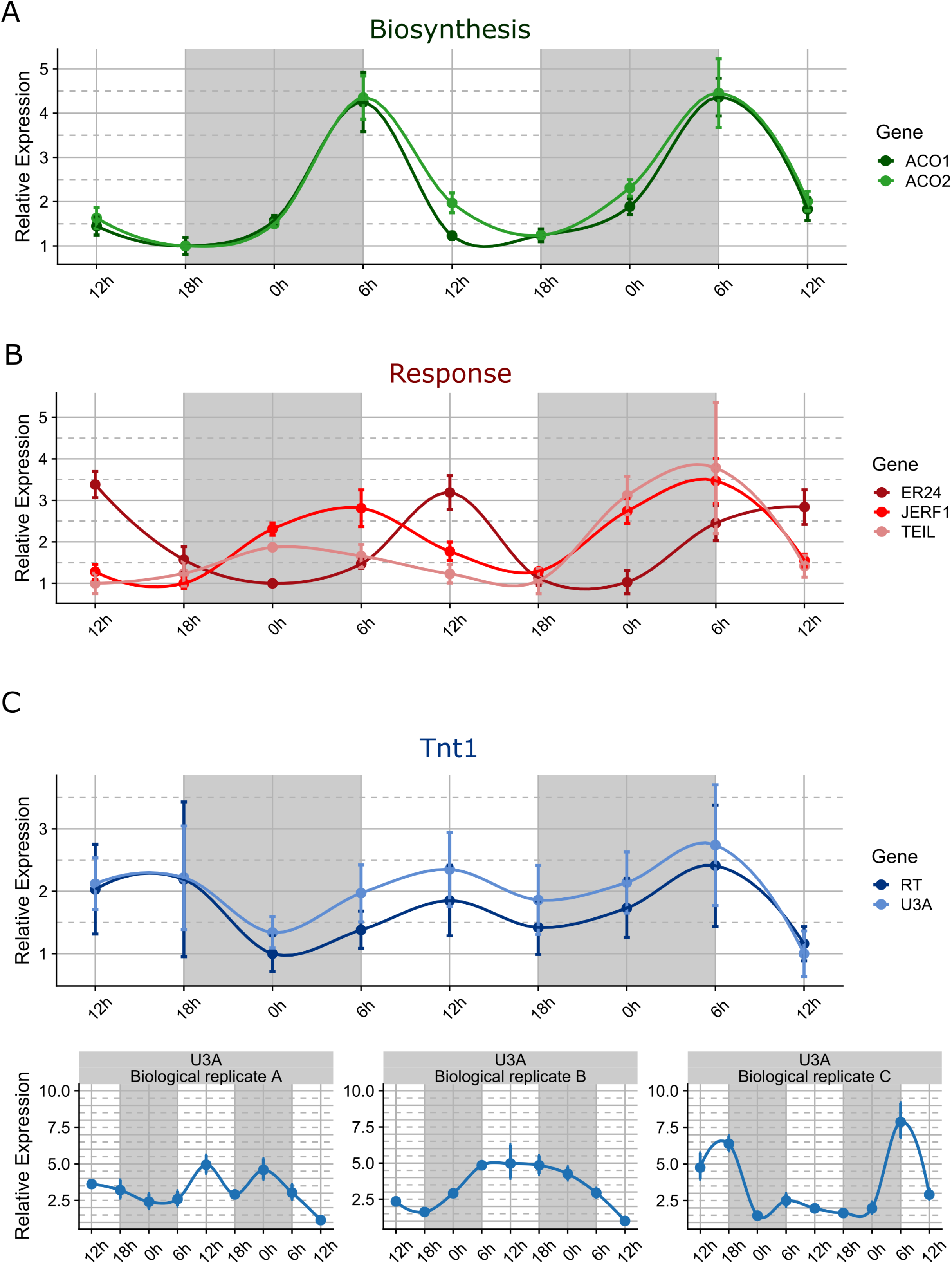
Relative expression of ethylene-related genes and Tnt1 in 15-days-old WT plants throughout 48 hours period. Plants were grown under a 12 hours light / 12 hours dark regime. Gray areas indicate dark periods. The lowest expression value for each gene was set to one. Bars represent standard deviation between three biological replicates, except for Tnt1 U3A, for which bars represent the standard deviation between three technical replicates.

To further analyze the connection between Tnt1 expression and ethylene biosynthetic and responsive genes, WT and HP plants (the same used for measurement of ethylene in gas chromatography) were also used for qRT-PCR experiments. Most genes assayed related to ethylene synthesis and response, were downregulated in HP plants when compared to the WT (Figure 3). Two copies of the ACC oxidase (*ACO1* and *ACO2*) gene, which participates in ethylene biosynthesis, were slightly downregulated (with relative expression ranging from 0.6 to 0.8 when compared to WT, except for *ACO2* in HP8 with expression 1.4 times higher than the WT line (Figure 3). The ethylene responsive genes *JERF1*, *ER24*, *SAR8.2b*, *TEIL* and *CHN48* were also downregulated in HP lines, except for *TEIL* in HP8, which had expression comparable to the WT (Figure 3). Finally, we also assayed three different Tnt1 coding domains (Gag, Integrase and Reverse Transcriptase), and they were downregulated in all HP plants (Figure 3).

**Figure 3.**
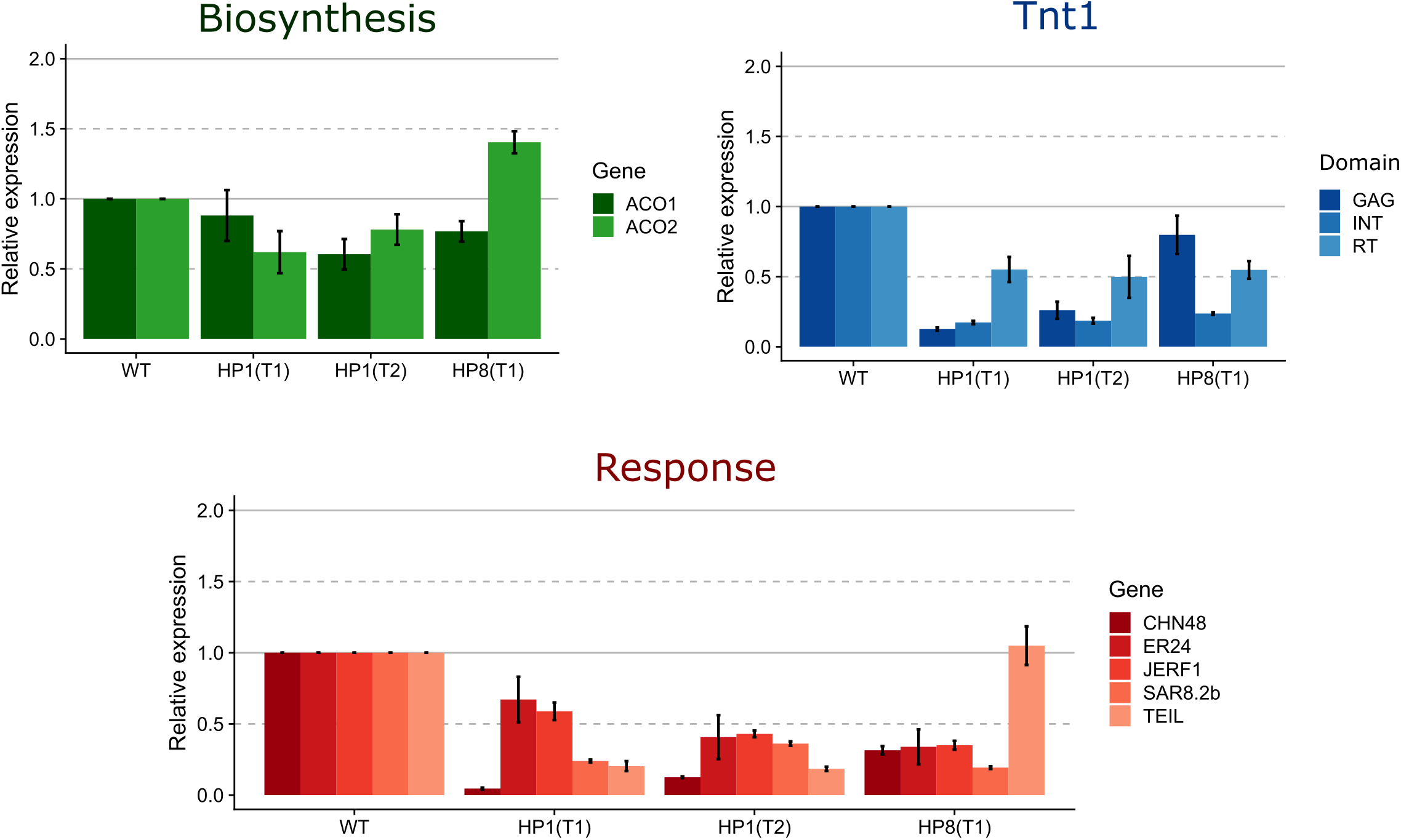
Relative expression of ethylene-related genes and Tnt1 in 90-day-old WT and HP plants. The expression level of WT was set as one for each gene. Values are linear average, with bars showing standard errors. Leaves used to extract the RNA for the quantitative PCRs were from the same plants used for the ethylene gas chromatography (Figure 1A).

### HP lines transcriptome profile differs from wild-type tobacco

Given the perturbations found in the ethylene biosynthesis and response pathways of HP lines, we asked whether these alterations could influence global gene expression. Thus, a whole transcriptome RNA-seq was performed on leaves of 45-day-old WT, empty-vector control plants (transformed with a vector with the same backbone but lacking the hairpin construction) and HP lines [HP1 (T1), HP1 (T2) and HP8 (T1)]. Because the most recent published tobacco genome is not yet complete, but is partly assembled into chromosomes by optical mapping, we expect a number of undefined nucleotides between scaffolds and is likely missing a number of gene annotations. Thus, we followed two approaches to process our RNA-seq data: (1) mapping reads against the tobacco whole genome in Solgenomics database and (2) mapping reads against unigenes from the tobacco database in Genbank.

Of the 35,519 annotated gene models in the tobacco genome, 16,331 had detectable expression levels [counts per million (CPM) > 1 in at least eight sequenced samples (including replicates)]. From these, we identified 932 differentially expressed genes (DEGs) in HP lines compared to the WT and control lines (FDR < 0.001, fold change ≥ 2). Hierarchical clustering of expression values for the 932 DEGs grouped samples in two classes, segregating HP lines from WT and control lines (Figure 4).

**Figure 4.**
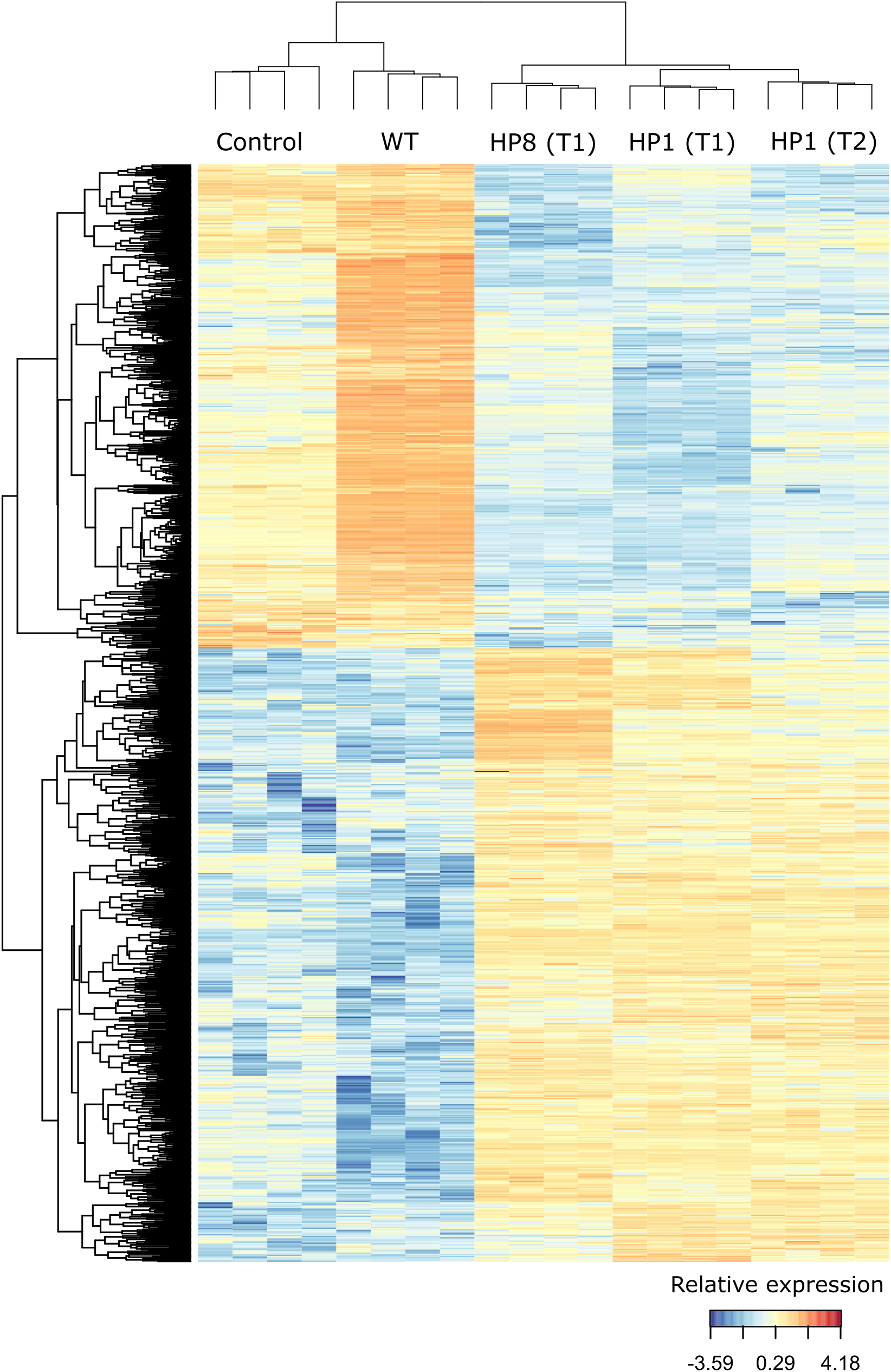
Hierarchical clustering analysis of 932 differentially expressed genes across different tobacco lines. Each column represents a biological replicate. Control line was transformed with a vector containing only the resistance gene.

Enrichment of Gene Ontology (GO) terms (p-value ≤ 0.01) identified a number of modulated biological processes in the HP lines (Figure 5). The most represented terms among the upregulated genes were: organophosphate metabolic process (GO:0019637), carbohydrate catabolic process (GO:0016052) and anion transport (GO:0006820); while metabolic process (GO:0008152), cellular process (GO:0009987) and cellular metabolic process (GO:0044237) were the most frequent terms of downregulated genes. Interestingly, we observed enrichment for stress-related processes in both sets of genes [upregulated: reactive oxygen species metabolic process (GO:0072593), defense response (GO:0006952), response to ethylene (GO:0009723) and reactive nitrogen species metabolic process (GO:2001057); downregulated: homeostatic process (GO:0042592), cellular homeostasis (GO:0019725) and cell redox homeostasis (GO:0045454)].

**Figure 5.**
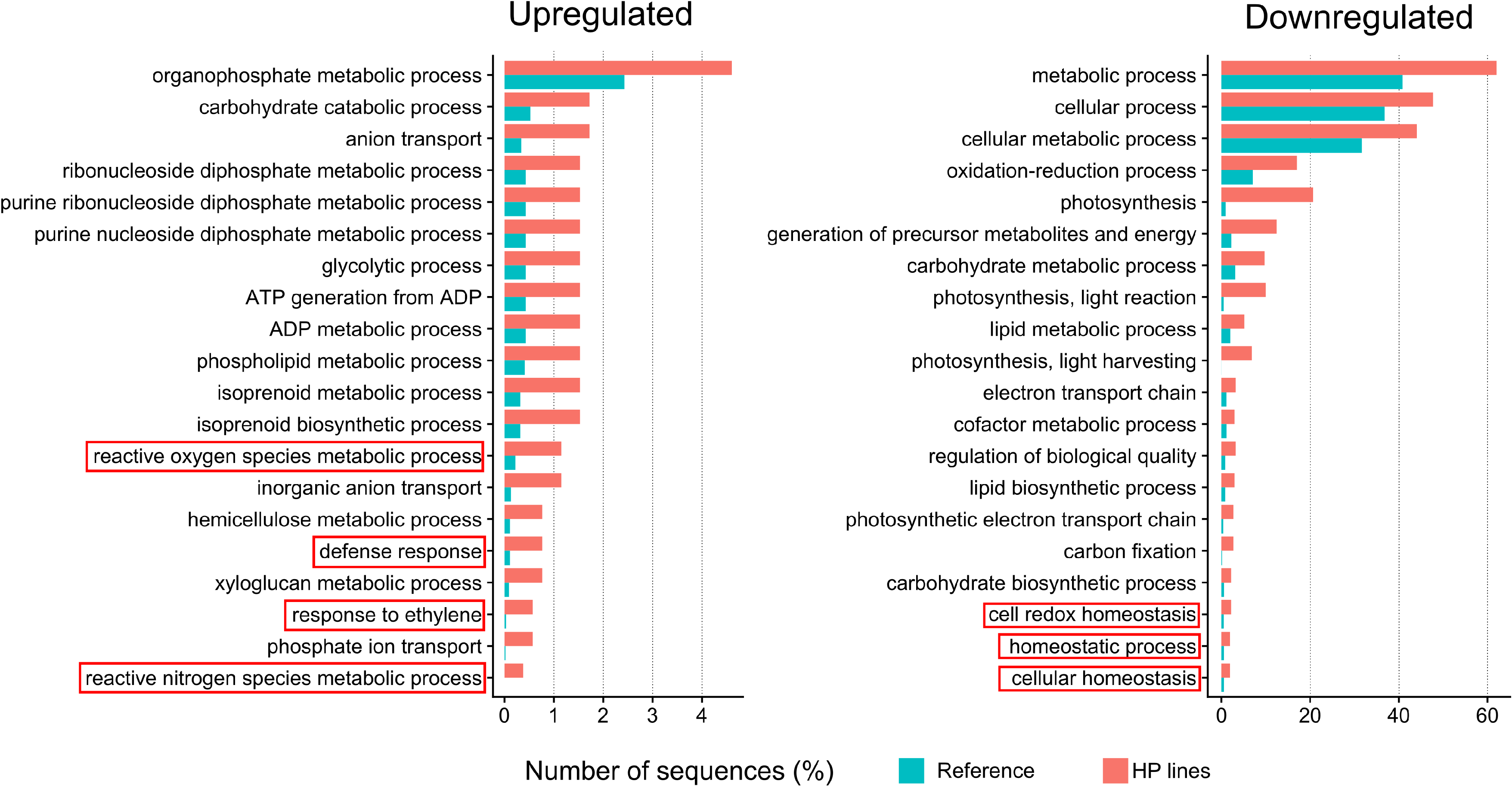
Gene ontology (GO) enrichment analysis (p-value ≤ 0.01) of the 932 differentially expressed genes between wild-type and hairpin lines. Only the 20 most representative biological processes are shown. Red boxes highlight processes related to stress response and homeostasis maintenance. The reference includes all expressed genes in all sequenced samples.

A second, more stringent approach was performed, in which each transgenic line (Control and HPs) were compared against WT. We then considered only DEGs consistently found in all HP samples and filtering out DEGs identified between WT and Control lines. Ninety seven DEGs were found (p-value ≤ 0.001, fold change ≥ 2), reflecting significant changes in gene expression. GO categorization of these 97 DEGs corroborates the results found in the previous analysis, with terms like “defense response”, “response to biotic stimulus” and “response to ethylene” appearing in the upregulated gene set (Additional file 1: Figure S2B). We used these 97 DEGs as seeds to start the inference of the gene regulatory networks using entropy based approach from gene expression patterns. The inference process was performed by selecting the predictors for each seed gene on each step. Thirty-five networks modules were identified revealing gene circuits in HP plants that were not identified in WT. From these, the most representative network module and which had the highest number of nodes connecting 44 genes is presented in Additional file 1: Figure S3.

### Phenotypic changes in HP lines

HP plants displayed a varied range of phenotypes. Lesion mimicry was readily observed in leaves of all four HP lines selected for this study (Figure 6A). These necrotic spots were present in T1 plants growing under normal conditions, but also in subsequent generations (up to T5) when exposed to stress, such as transfer from *in vitro* cultivation to soil (Figure 6A).

**Figure 6.**
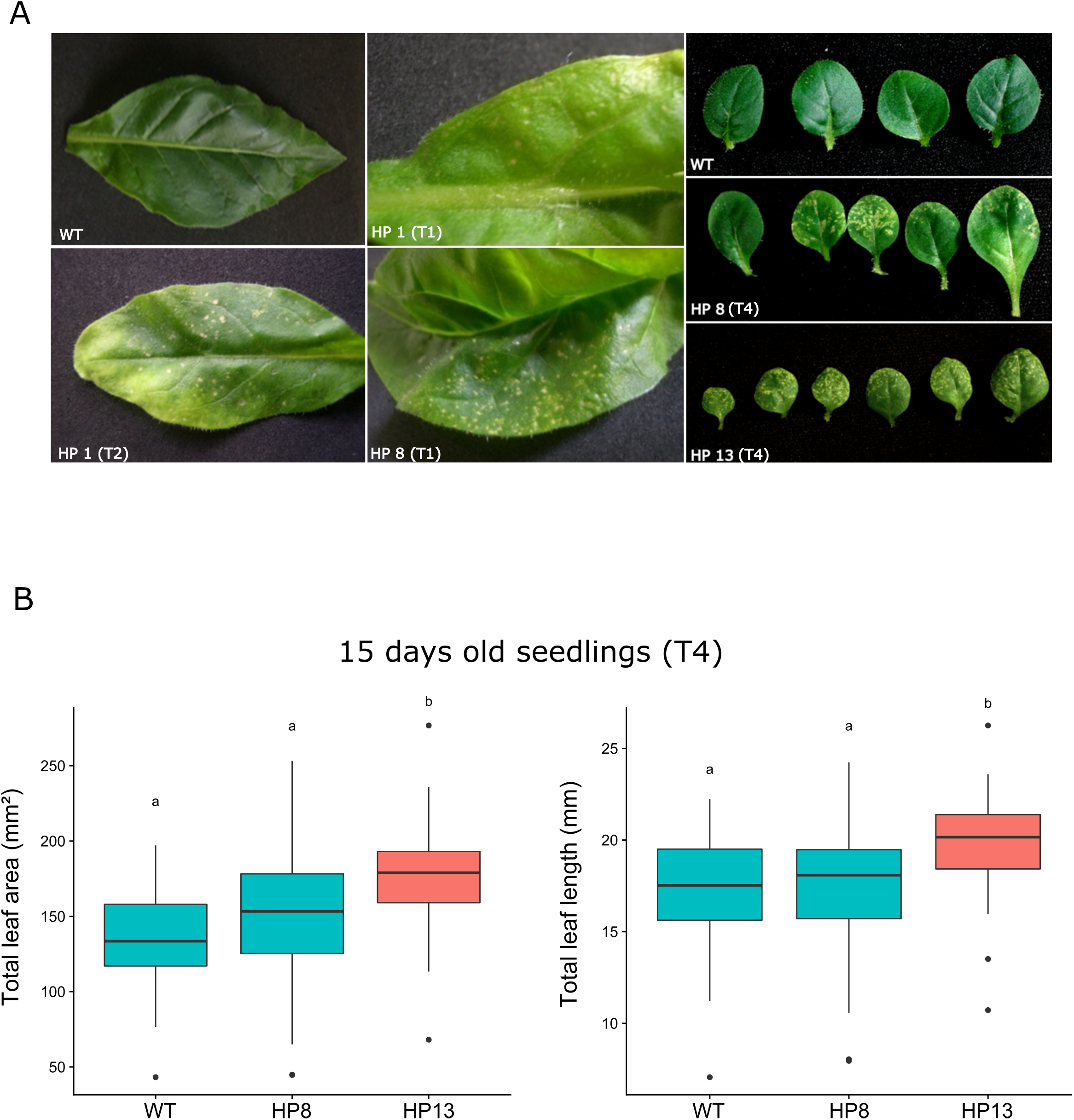
Phenotypes of aerial parts in transgenic RNAi lines. **(A)** Leaves of wild-type (WT) and transgenic (HP) *Nicotiana tabacum* plants growing *in vitro*. Wild type presents normal leaves while HP lineages present cell death spots. Left panel (WT, T1 and T2) displays leaves of 45-days-old plants used for the transcriptome sequencing. **(B)** Total leaf area and total leaf length (measured from the first leaf pair) of 15-days-old seedlings. Letters represent statistically significant differences between lineages.

To explore the consequences of interfering with Tnt1 expression we compared the organization of the shoot and root, as well as the germination rate of WT and HP lines (HP8 and HP13). Two measures were considered at two-leaf stage plants (15-days old): total foliar area and the maximum distance across the longer longitudinal axis of the two leaves. Under these parameters, the HP13 line showed significantly increased growth of the shoot system; the total foliar area was 30% greater than the wild type and 15% longer in the longitudinal axis (Figure 6B, Table 1).

**Table 1.** Leaf size comparison between 15 days old seedlings of WT and HP lines (T4).

| Line | Total leaf area (mm<sup>2</sup>) | Total leaf length (mm) |
| --- | --- | --- |
| WT | 135.26 ± 31.86<sup>a</sup> | 17.20 ± 3.14<sup>a</sup> |
| HP8 | 147.05 ± 44.41<sup>a</sup> | 17.27 ± 3.51<sup>a</sup> |
| HP13 | 175.86 ± 34.71<sup>b</sup> | 19.75 ± 2.68<sup>b</sup> |

Letters indicate statistically significant differences (p-value ≤ 0.05). Total leaf length was measured as the maximum distance across the longer longitudinal axis of the two leaves.

Several morphological phenotypes were observed in roots. HP lines growing *in vitro* developed longer root hairs close to the root tip when compared to WT (Figure 7A, Additional file 1: Figure S4A). Root growth was also altered in HP lines: primary root length, surface and volume were significantly smaller in 15-days-old plants (Figure 7B; Table 2). The same result was observed when comparing the whole root system (Additional file 1: Figure S4B; Table 2). HP lines also tended to produce fewer lateral roots (Figure 7B; Additional file: Figure S4B; Table 2).

**Figure 7.**
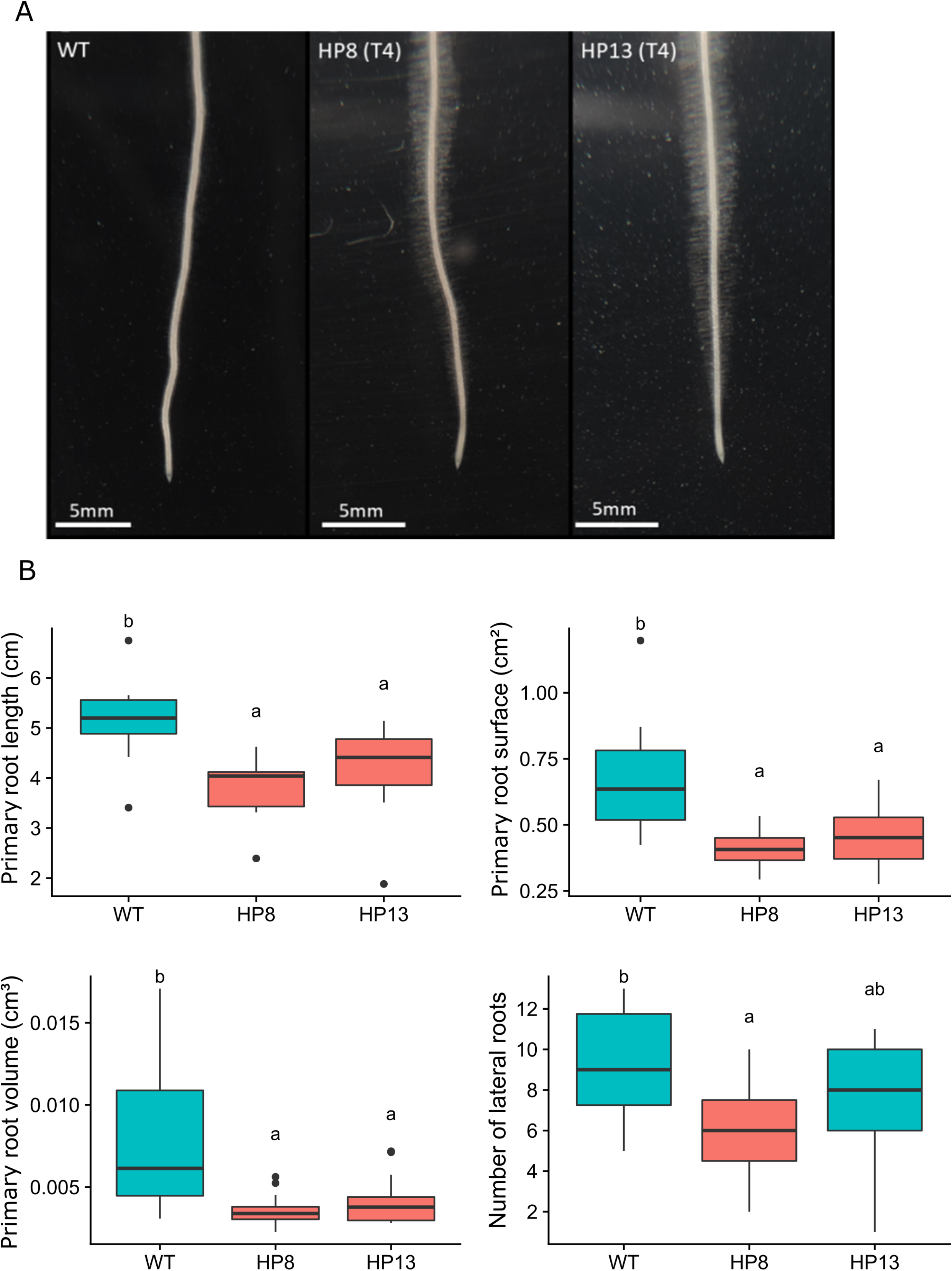
Phenotypes of the root system in transgenic RNAi lines. **(A)** Primary root of wild-type (WT) and transgenic (HP) lines. Root hair production is increased in HP lines. **(B)** Comparison of morphological parameters in the primary roots of WT and HP 15-days-old seedlings. Letters represent statistically significant differences between lineages.

**Table 2.** Comparison between the root system of 15 days old WT and HP seedlings.

| Sample | Line | Length (cm) | Surface (cm<sup>2</sup>) | Volume (cm<sup>3</sup>) | Number of lateral roots |
| --- | --- | --- | --- | --- | --- |
| Primary root | WT | 5.14 ± 0.87<sup>b</sup> | 0.68 ± 0.24<sup>b</sup> | 0.0079 ± 0.0044<sup>b</sup> | 9.30 ± 2.67<sup>b</sup> |
|  | HP8 | 3.83 ± 0.58<sup>a</sup> | 0.40 ± 0.07<sup>a</sup> | 0.0036 ± 0.0009<sup>a</sup> | 6.00 ± 2.54<sup>a</sup> |
|  | HP13 | 4.22 ± 0.88<sup>a</sup> | 0.46 ± 0.12<sup>a</sup> | 0.0042 ± 0.0015<sup>a</sup> | 7.77 ± 2.86<sup>ab</sup> |
| Total root system | WT | 11.36 ± 3.65<sup>b</sup> | 1.34 ± 0.52<sup>b</sup> | 0.0140 ± 0.0078<sup>b</sup> | NA |
|  | HP8 | 7.49 ± 2.31<sup>a</sup> | 0.73 ± 0.24<sup>a</sup> | 0.0060 ± 0.0025<sup>a</sup> | NA |
|  | HP13 | 8.72 ± 2.89<sup>ab</sup> | 0.92 ± 0.40<sup>a</sup> | 0.0082 ± 0.0047<sup>a</sup> | NA |

NA – Not applicable. Letters indicate statistically significant differences (p-value ≤ 0.05).

We also compared germination of 6-year-old and fresh seeds of WT and HP lines considering two parameters: root and cotyledon emergence. Germination was delayed in HP seeds when compared to the WT (Figure 8; Additional file 1: Figure S5). For example, at nine days after sowing most of the fresh WT seeds had both root and cotyledons emerged, with only 1.33% of the seeds partially germinated with emerged roots but no cotyledon. On the other hand, in HP8 and HP13 lines 21.33% and 10.67% seedlings were partially germinated, respectively, for the same period (Figure 8). HP13 line has the most impacted germination rate (as measured 15 days after sowing), 58.92% compared to 94.66% in WT plants (Table 3). Not only the germination rate was altered, but a decrease in seed vigor was indicated by both mean germination time (MGT) and germination speed index (GSI) (Additional file 1: Figure S5; Table 3). Although HP lines had a slight reduction in germination rate compared to the WT (89.33% in HP13 versus 98.67% in WT; Additional file 1: Figure S5; Table 3), only the MGT was significantly different in HP lines (Additional file 1: Figure S5; Table 3).

**Figure 8.**
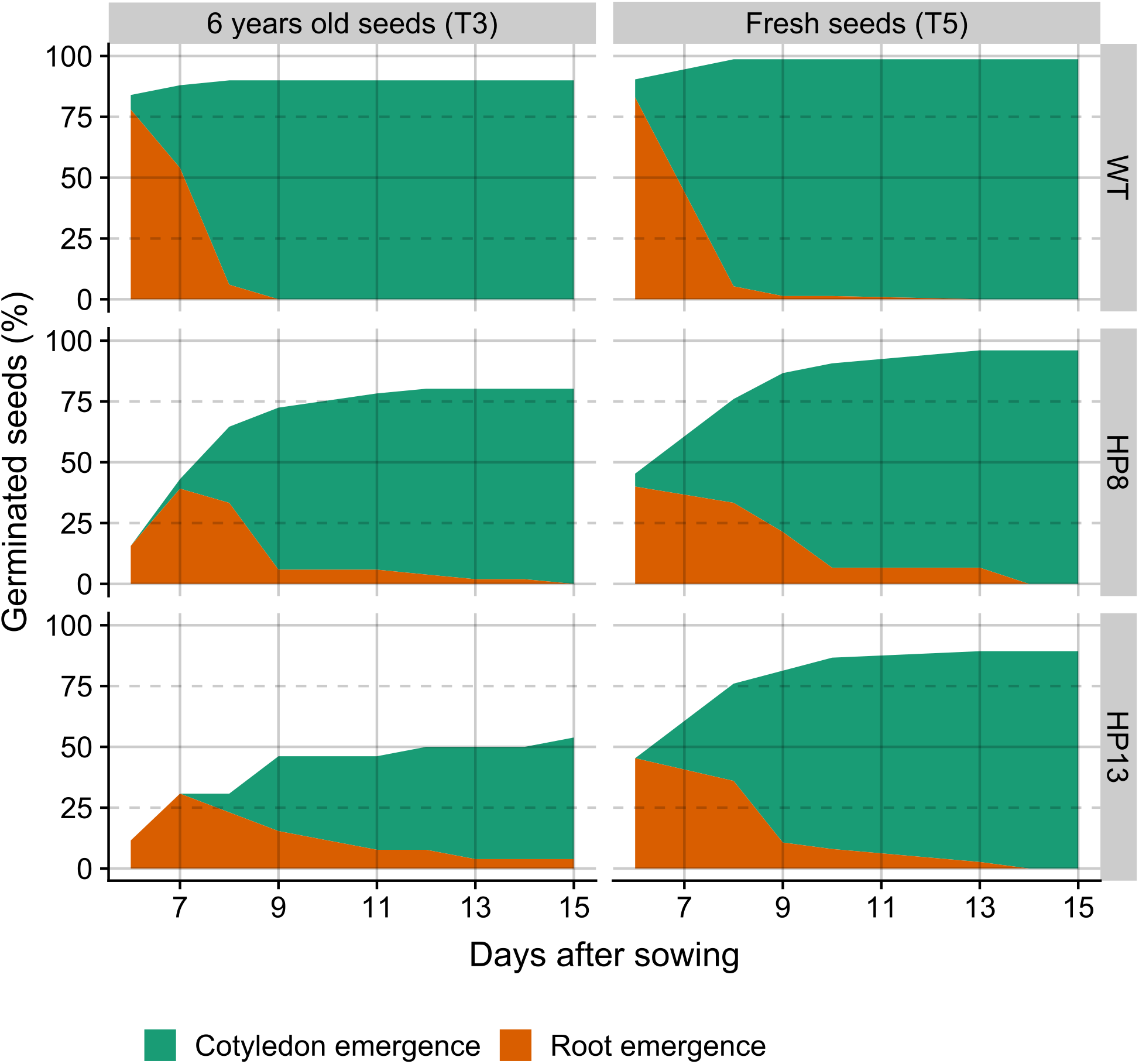
Germination progress in wild-type (WT) and transgenic RNAi (HP) lines. Seeds were observed from 6 to 15 days after sowing. Orange area represents the percentage of seeds displaying root emergence, while green denotes seedlings in which both root and cotyledon emergence had occurred.

**Table 3.** Germination comparison between fresh and six years old seeds from WT and HP lines.

| Seed age | Line | Mean germination time (days) | Germination speed index | Germination rate (%) |
| --- | --- | --- | --- | --- |
| Six years old | WT | 5.68 ± 0.45<sup>c</sup> | 4.26 ± 0.64<sup>c</sup> | 94.67 ± 9.24<sup>b</sup> |
|  | HP1 | 6.79 ± 0.19<sup>bc</sup> | 3.46 ± 0.30<sup>bc</sup> | 89.33 ± 4.62<sup>b</sup> |
|  | HP5 | 9.26 ± 1.33<sup>a</sup> | 2.52 ± 0.21<sup>a</sup> | 89.28 ± 6.02<sup>b</sup> |
|  | HP8 | 7.35 ± 0.44<sup>abc</sup> | 3.07 ± 0.55<sup>abc</sup> | 85.49 ± 12.27<sup>b</sup> |
|  | HP13 | 8.53 ± 0.04<sup>ab</sup> | 2.00 ± 0.28<sup>ab</sup> | 58.93 ± 7.18<sup>a</sup> |
| Fresh | WT | 6.17 ± 0.07<sup>b</sup> | 3.81 ± 0.31<sup>b</sup> | 98.67 ± 2.31<sup>a</sup> |
|  | HP8 | 7.53 ± 0.25<sup>a</sup> | 3.35 ± 0.15<sup>a</sup> | 96.00 ± 4.00<sup>a</sup> |
|  | HP13 | 7.33 ± 0.38<sup>a</sup> | 3.18 ± 0.35<sup>a</sup> | 89.33 ± 6.11<sup>a</sup> |

Letters indicate statistically significant differences (p-value ≤ 0.05).

### Tnt1 genomic insertions

We asked how interfering with Tnt1 expression could result in the phenotypes and transcriptome profile changes observed in HP lines. The presence of Tnt1 insertions in the vicinity of genes and / or within introns could lead to an indirect reduction in mRNA of those genes in HP lines via the RNAi mechanism. To avoid this bias, we first identified genomic copies of Tnt1 by searching and annotating its distinct domains (i.e. U3A, U3B, U3C, U5, the whole LTR, as well as the coding domains GAG, PROT, INT, RT and RNH).

A total of 276 U3A, 155 U3B and 35 U3C regions were found, each of them being part of a complete Tnt1 insertion, an incomplete insertion or even a solo LTR. Not surprisingly, many Tnt1 insertions are found in close proximity to scaffold borders, making it impossible to determine the completeness of most of the insertions. Next we identified the first gene present within a 5kb distance (in both upstream and downstream direction) of each U3 region. From the 216 genes found near Tnt1 insertions, only 11 were differentially expressed, being 8 upregulated and 3 downregulated. Interestingly, 127 of these 216 genes had no detectable expression levels in our RNA-seq experiment, suggesting that Tnt1 insertions may play a role in silencing mechanisms of neighbor genes.

Finally, given that there is not yet a complete assembly of the tobacco genome, it is important to note that these results are possibly an underrepresentation of the total number of genomic Tnt1 insertions. As described above, because of their repetitive nature, Tnt1 sequences are prone to appear next to scaffold borders, and a number of unknown insertions are expected to exist in the gaps between scaffolds.

### Tnt1 LTR as a putative source of sRNAs harboring GCC-motifs

Because TEs are known to be a source of noncoding RNAs, we checked if Tnt1 transcripts could form secondary structures. Thus, we performed a folding prediction of part of the Tnt1A mRNA, which includes three 3’ U3 GCC-like motifs (Figure 9). The prediction reveals stem-loop secondary structures, and GCC-like motifs would be located in the arms of the putative hairpin loops (Figure 9), with the folded RNA free energy of dG = −24 kcal/mol.

**Figure 9.**
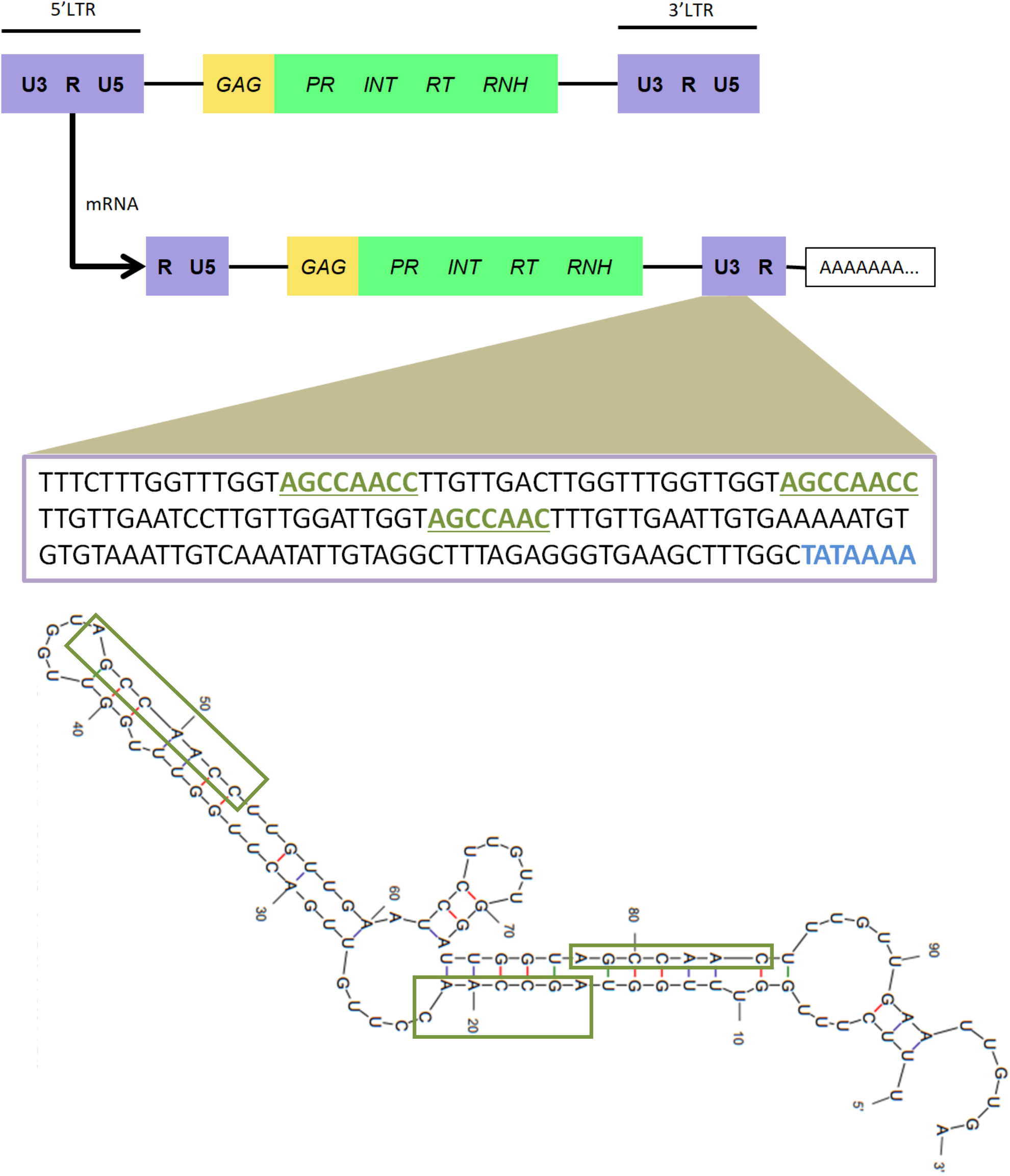
The discovery of putative Tnt1-derived GCC-box small RNAs in the promoter region of Tnt1A. Tnt1 retrotransposon genomic and transcript structures. The text box bellow the Tnt1 scheme represents the 3’ U3 region sequence of Tnt1A. Green underlined words indicate the 3 GCC-like motifs and TATA box is in blue. Bottom figure depicts the folding prediction of a 98 bp part of the Tnt1A mRNA sequence containing the three GCC-like boxes (green rectangles).

Next, we used PsRNATarget (see “Methods” section for details) to search for potential targets of three putative sRNAs derived from the stem-loop secondary structures predicted in the previous step, each of them containing one GCC-like motif. To account for both transcriptional and post-transcriptional possible targets of sRNAs, searches were performed in two datasets: (1) a PsRNATarget built-in library of 25,398 *Nicotiana tabacum* SGN unigenes, in which only the coding sequences are considered, and; (2) sequences of 3kb upstream of the 932 DEGs identified in this work, aiming to understand if the promoter region of such genes could be potential targets of the sRNAs. Because GCC motifs are regulatory sequences, one would expect them to be found in the promoter region of genes, particularly among the DEGs found in the present study.

As results, we found 224 putative targets within the coding sequences of the SGN unigenes library, from which 14 were present in our set of 932 DEGs. When targeting the upstream 3kb region, a total of 106 out of the 932 DEGs showed potential targets for the putative sRNAs, of which 67 were upregulated and 39 downregulated in HP lines. Based on these results, our hypothesis is that Tnt1 exerts transcriptional regulation of some genes via sRNAs.

## DISCUSSION

### The dynamics of Tnt1 expression in WT tobacco

Expression of the Tnt1 retroelement is known to be induced by both biotic and abiotic factors. Protoplast preparation by fungal extracts is the most studied way to induce Tnt1 expression, and was used to demonstrate mobilization of the element [32]. Likewise, expression of Tnt1 was reported to be induced upon viral infection and seems to be linked to plant defense responses, although its role in such cases is still unclear [33]. Interestingly, expression of Tnt1 subfamilies (Tnt1A, Tnt1B and Tnt1C) is induced by different stress-associated molecules, with Tnt1A being particularly upregulated by cryptogein and methyl jasmonate [29]. Activation of different subfamilies is accredited to the presence of specific promoter motifs in each of their U3 regions (U3A, U3B and U3C). Although Tnt1A responsiveness to ethylene has been previously hypothesized due the presence of repetitive GCC-like motifs in its U3A region [31], no experimental data was available to support this idea. Our results demonstrate that after exposure to the gaseous-hormone ethylene, expression of Tnt1 is upregulated (30 to 40-fold increase).

Expression of ethylene biosynthetic and responsive genes follow a basal circadian cycle, however that does not mean senescence, stress-responses, and other mechanisms dictated by ethylene also oscillates in a circadian fashion. Results from an experiment designed to address if Tnt1 followed either of the two ethylene expression patterns (circadian associated or not) is consistent with the non-circadian response and support basal and continuous Tnt1 expression. The hypothesis raised by the results presented is that Tnt1 basal expression is not controlled by ethylene, but rather induced by the hormone only under specific conditions, probably requiring additional signaling or an elevated ethylene concentration.

Likewise, expression of Tnt1A, as indicated by quantification of both the RT domain and the specific U3A region, oscillate in a non-circadian fashion and greatly differs between biological replicates growing under the same controlled conditions. These results suggest that, under normal conditions, Tnt1A expression is not dictated by ethylene but rather by unknown, possibly stochastic, factors. In this scenario, ethylene induction of Tnt1A expression seems to be subject of additional regulation and may require exposure to a minimum threshold of the hormone.

### Perturbations in Tnt1 expression lead to phenotypic and transcriptomic changes

Since Tnt1 has basal expression in different tissues [20], we sought to understand whether its suppression would culminate in detectable phenotypes. Thus, we generated transgenic RNAi lines (herein referred as HP lines) expressing a hairpin targeting the RT domain of Tnt1. With this approach, Tnt1 transcripts were consistently downregulated upon ethylene treatment.

Different HP lines displayed recurring phenotypes, namely lesion mimicry in leaves, underdevelopment of the root system, overproduction of root hairs and early loss of seed viability. Most of these developmental aspects are regulated by ethylene to some extent. For instance, root hair production is induced by ethylene in *Arabidopsis* [34–36]. Interestingly, mutation in the *Arabidopsis RHD6* gene, which mediates ethylene-response during root hair formation, leads not only to a decrease in root hair production but also to a shift in their initiation site towards the root base [35], which is the opposite effect observed in HP lines.

In addition, the germination process is mediated by counteracting effects of ethylene and ABA in numerous species, including tobacco (see 37 for a review). It is possible that HP seeds lose viability via early partial release of dormancy by an overproduction of ethylene. Accordingly, we demonstrated that HP plants overproduce ethylene when compared to the WT.

Functional categorization of the 932 DEGs in HP lines reveals enrichment of stress-related processes among upregulated genes, while metabolic and homeostatic processes in the downregulated set of genes. This pattern is supported in a second and more strict analysis that pointed to 97 modulated genes. Among these DEG, several genes related to ethylene biosynthesis and plant defense response were upregulated in all HP lines but not in the WT as follows: two ACC oxidase/ethylene forming enzymes (EFEs), involved in the last step of ethylene synthesis (accession numbers AB012857 and Z29529)[38]; five pathogenesis-related protein (PR) family members (EH620111, EH621793, X03913, M29868 and X51426)[39–41]; phospholipase D which participates in signal transduction cascades in stress responses [42]; and an ethylene responsive gene induced during the pathogen-induced systemic acquired resistance, Sar8.2b (EH621848) [43]. In addition, we identified the repression of several chlorophyll a/b binding proteins in HP leaves. Senescence is marked by a decline of the photosynthetic apparatus and mobilization of nutrients from senescing leaves to growing tissues, culminating in cell death [44]. Likewise, hydroxy-methylglutaryl-coenzyme A reductase (HMGR)-like genes were up-regulated in HP plants. This enzyme participates in the steroid derivatives synthesis that follows pathogen infections [45]. This possibly corresponds to the spread of the initial stress signal throughout the plant. Although at this point it is not possible to distinguish between primary and secondary effects, we propose that the retrotransposon not only responds to some biotic and abiotic stresses, but also fine-tunes their occurrence and progression. Likewise, our network analysis strengthened the emergence of a new pattern of expression and gene regulation in the HP plants, and uncovered similar biological processes (e.g. defense and stress response) as the main changes in HP lines.

### Tnt1 acting as modulator of plant development and response to stress

Curiously, the expression kinetics differ between samples used for the RNA-seq experiment and those assayed by RT-qPCR, i.e. while ethylene-related genes were upregulated in the former, most of these genes were downregulated in HP lines used for qRT-PCR. Because samples used for qPCR were sealed in a container prior to harvesting, they were exposed to an increased concentration of the hormone. Accordingly, these samples showed downregulation of Tnt1 expression (GAG, INT and RT domains) in comparison to WT plants, much like the results obtained after application of exogenous ethylene (compare Figures 1A and 3). Thus, differently from WT, HP plants growing under normal conditions overproduce ethylene and behave as if constantly responding to stress, while showing a decreased response to stress when exposed to higher concentrations of ethylene. These results are consistent with Tnt1 modulation of two distinct processes: (1) maintaining homeostasis during normal development and (2) fine-tuning stress responses possibly mediated by ethylene.

Finally, we combine our results to propose a model of the interaction between expression of Tnt1, ethylene biosynthesis genes and ethylene responsive genes that contain GCC motifs in their promoter (Figure 10). In this model, we consider two different conditions: a period of time when a WT plant exists under ideal conditions for normal growth and another under a stress pressure. Under normal growth conditions, WT tobacco plants express Tnt1 in a basal level, which fluctuates within a range of up to 8-fold, as observed in our circadian experiment. Genes involved in ethylene biosynthesis and other ethylene responsive genes that have GCC motifs in their promoters are also expressed in a circadian fashion (since ethylene is required in various moments of plant development and not only during plant defense responses). Upon a stress stimulus, ethylene biosynthesis is increased and this is one of the events that define the commencement of the defense responses. The overproduction of ethylene triggers the upregulation of genes that contain GCC motifs in their promoter (such as Tnt1 and other responsive genes). According to our analysis, it is possible that Tnt1A mRNA can be a source of sRNAs that target GCC motifs in the promoter region of genes, or alternatively, in the 3’UTR of mRNAs. Thus, as Tnt1 is upregulated by ethylene, there is a turning point in which the Tnt1A sRNA production and consequent transcriptional inhibition of Tnt1 and target genes overcomes the ethylene induction, thus lowering the amount of mRNA of Tnt1 and of ethylene responsive genes. After ethylene responsive genes reach their maximum of expression, right before Tnt1A mRNA-derived sRNAs start to inhibit their transcription, it is likely that the defense responses have taken place and were sufficient to overcome the initial stress, thus removing this stimulus and lowering the expression of ethylene biosynthetic genes through ethylene auto-inhibition. The decrease in ethylene production removes the induction signal for Tnt1 transcription, lowering Tnt1 transcription and thus lowering Tnt1A sRNA production also. This way the system is pushed back to its “normality” after the stress has been overcome.

**Figure 10.**
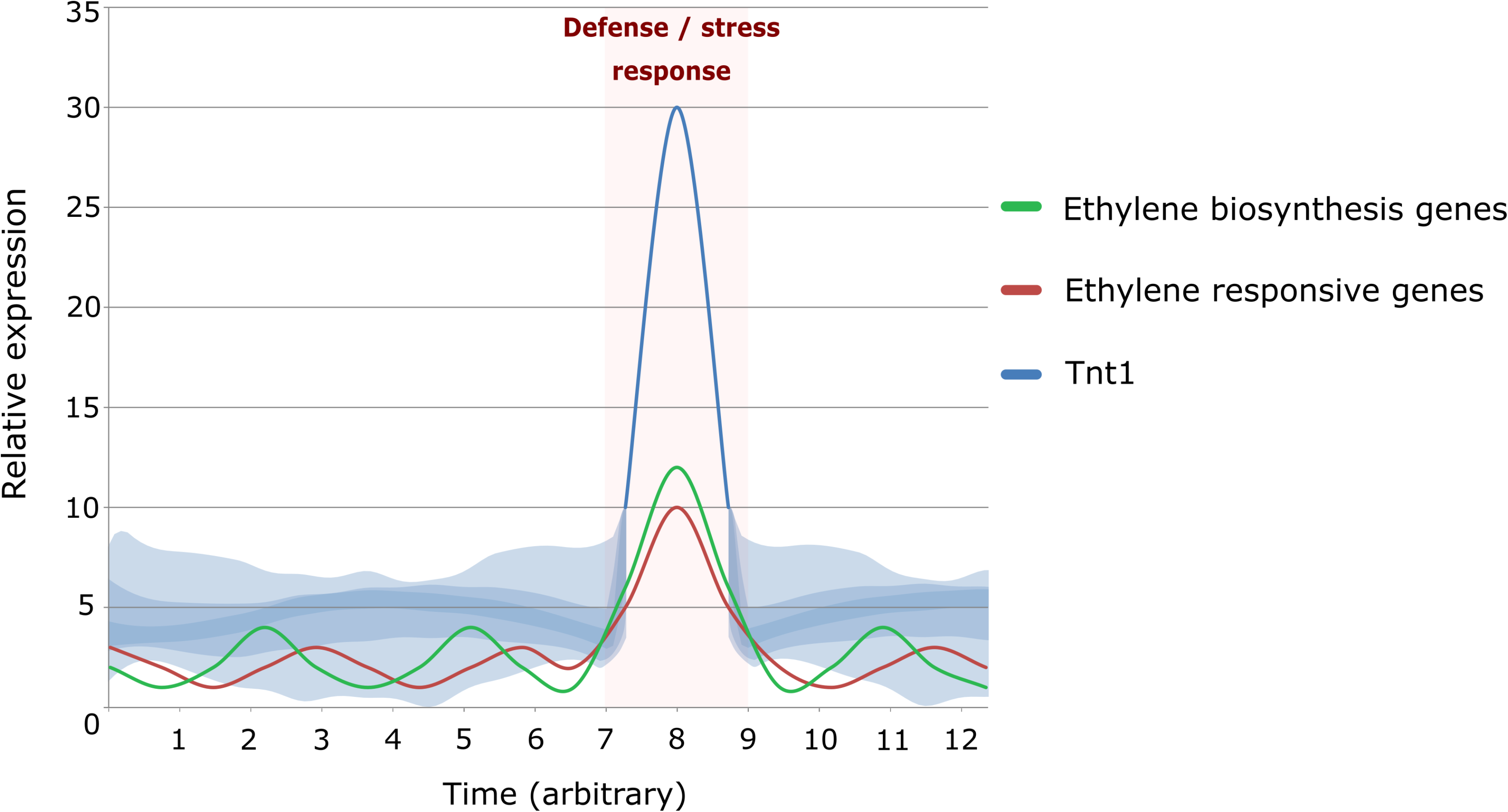
Model proposing a dynamic equilibrium between expression of defense response genes and Tnt1 in WT tobacco. Expression values are based on our observations, except for ethylene biosynthesis and responsive genes during defense/stress response, for which values are hypothetical. During normal development (T0 – T7), ethylene biosynthetic/responsive genes are expressed in a circadian fashion, while Tnt1 fluctuates within basal range with no clearly delimited periodicity (depicted as a blue cloud). Upon stress stimulus, ethylene biosynthesis is upregulated, which also induces expression of genes that contain GCC motifs in their promoter, including Tnt1A (T8). Tnt1-U3A GCC-like motifs sRNAs overcomes ethylene induction and promotes transcriptional inhibition of GCC motifs in other ethylene-responsive gene promoters (T8 – T9). Expression of Tnt1 and ethylene-responsive genes returns to normal levels due to the repression mediated by its own sRNA (T9 – T12).

## CONCLUSIONS

Although transcriptionally active TEs are commonly taken as a potential threat to their host organisms, there are recent reports of other TEs playing important roles for plant development. Such is the case of the MIKKI retrotransposon modulating root development in rice [19]. Another interesting case is found in *Solanum* species, in which the MESSI retrotransposon family is expressed specifically in the shoot apical meristem, suggesting that these TEs can respond to developmental signals [46]. Our findings suggest that active retrotransposons, not only domesticated transposable elements, can play a significant functional role in their host organisms. We hypothesize that Tnt1 can exert transcriptional control over itself as well as other endogenous genes. In our model we propose a potential new biological role for Tnt1. Upon stress induction, Tnt1 would provide feedback control to ethylene-mediated gene regulation in tobacco defense responses, bringing the system back to a homeostatic condition after the initial stress stimulus has been overcome. Further studies on the progression of Tnt1 expression during stress response and recovery, as well as small RNA-seq experiments, can validate and bring new insights to the model presented herein.

## METHODS

### Plant samples and genetic transformation

Plants of *Nicotiana tabacum* cv Xanthi XHFD8 were used for genetic transformation to produce transgenic RNAi lines, herein referred as “HP lines”. We design a hairpin construct aiming to target a conserved region of the Tnt1 RT domain, shared by the Tnt1A, Tnt1B and Tnt1C subfamilies. To achieve the hairpin construction, a 273bp fragment of the Tnt1 (accession number X13777.1) reverse transcriptase was amplified from *N. tabacum* genomic DNA (forward primer 5’ CGGGATCCATCTCAGCAGAAGTACAT 3’, reverse primer 5’ CCATCGATACTTCCCAATGTTCC 3’). This fragment was cloned in the expression vector pHANNIBAL (accession number AJ311872.1) in both sense and antisense directions, separated by an intron, in order to express a Tnt1 hairpin. Expression cassettes were excised from pHANNIBAL and transferred to the binary vector pCAMBIA1201, generating pCAMBIA-Tnt1-RT. Control plants were transformed only with the pCAMBIA1201 backbone containing the hygromycin resistance gene. *Nicotiana tabacum* foliar discs were transformed with pCAMBIA-Tnt1-RT through *Agrobacterium tumefaciens* (LBA4404) co-culture, according to a previously established method [47]. Seventeen transgenic HP plants were generated with this cassette. Regenerated *in vitro* transgenic plants were cultivated in MS20 media with the proper antibiotic, under a 14-hour photoperiod at 24°C.

### Phenotyping of transgenic lines

For all phenotyping experiments, seeds of WT and HP lines were first primed (24 hours in sterile distilled water at 10°C) and then germinated and grown *in vitro* in MS20 medium [48] under a 14-hour photoperiod at 24°C. Comparison of leaf area and leaf length was done using pictures taken from 15-day-old seedlings and then used for measurements with Fiji distribution of ImageJ [49]. For root analyzes, plants were grown in 45° inclined Petri dishes. Pictures were taken from plants 15 days after sowing. These pictures were used for measurements with the ImageJ SmartRoot plugin [50]. Germination assay was performed through daily observation of root and cotyledon emergence. Germination Speed Index (GSI) and Mean Germination Time (MGT) were calculated following [51]. All experiments were done using three replicates of 25 plants per line.

### RNA-seq and reads processing

Total RNA from leaves of 45-day-old plants were frozen in liquid nitrogen. Four biological replicates from three independent HP lines, one Control and one WT were harvested. TRI Reagent® (Sigma-Aldrich) was used for RNA isolation according to the manufacturer’s instructions. Samples were treated with DNaseI (Ambion) and ribosomal RNA was depleted using Ribominus Plant Kit (Invitrogen) following the manufacturer’s instructions. The cDNA libraries were made using SOLID Total RNA-seq Kit (Ambion), according to Whole Transcriptome Library Preparation for SOLID Sequencing Protocol (Life Technologies). The handling of the beads for sequencing was done strictly according to the SOLID 3 System Templated Bead Preparation Guide (Life Technologies). The run of the samples followed the SOLID 3 System Instrument Operation Guide (Life Technologies). The RNA-seq produced a total of 602,744,341 single-end reads (GenBank GEO accession GSE44027), which were trimmed and quality filtered using Trimmomatic [52] with default settings.

### Bioinformatics and gene expression analysis of RNA-seq data

The RNA-seq reads were first mapped against the tobacco genomic assemble [53] using HISAT2 [54] set to default parameters. Differentially expressed genes (DEGs) were defined using edgeR [55]. Lowly expressed genes were filtered out based on a minimum of 10 counts per million in at least eight sequenced samples (including replicates). Sample normalization was performed using the trimmed mean of M-values (TMM) method. Threshold for DEGs was set using a false discovery rate (FDR) of < 0.001, yielding 932 genes. In a more stringent approach, reads were also mapped against 24,069 unigenes from the tobacco database in Genbank. The normalization of transcriptome data was done based on RPKM expression measure [56], square root, and Bonferroni’s correction. Using statistical t-test with p-value < 0.001 we generated two subsets of modulated genes, the first comparing the three HP in contrast to WT lines, and only the modulated in the three HP were considered. The second subset comprises the genes modulated comparing HPs with Control, in order to filter out genes possibly modulated due to the transgenesis process. This method identified 97 DEGs. Gene ontology (GO) analysis was performed using Blast2GO [57]. Enrichment tests among upregulated and downregulated genes were made comparing the set of expressed genes in all samples with those expressed in HP lines with p-value ≤ 0.01.

### Gene regulatory networks

Connections between genes were inferred by adopting the mean conditional entropy (MCE) from the observed gene expression data [58]. The MCE is an information measure that for each target gene indicates the contribution of its predictors to correctly detect the target behavior in a multivariate way.

### Annotation of Tnt1 genomic insertions

Sequences for each domain of Tnt1A (X13777), as well as known variant sequences of U3A (AJ227998, AJ228000, AJ228002 – AJ228006, AJ228008, AJ228010 – AJ228012, AJ228014 – AJ228017), U3B (AJ227999, AJ228007, AJ228009, AJ228013) and U3C (AJ228001) were imported into Geneious Prime 2020.0.5 software (https://www.geneious.com). Next the “Live Predict & Annotate” tool (which performs a blast-like search) were used with a threshold of 60% to find and annotate each Tnt1 domain in the tobacco genome [53]. The first gene present within a 5kb distance (both upstream and downstream) of each insertion was retrieved, as well as information about its orientation in relation to the insertion (sense or antisense).

### Small RNAs target prediction

Prediction of sRNAs putative targets were performed using the PsRNATarget tool [59] with default parameters. For target sites within coding sequences, the build-in *Nicotiana tabacum* SGN unigene cDNA library was used. Promoter region of the differentially expressed genes were extracted from the tobacco genome latest release [53], considering 3kb upstream of each gene.

### Ethylene treatment and ethylene gas chromatography

Tnt1 induction by ethylene was performed by placing plants in sealed containers and then ethylene was taken from a concentrated stock (Alltech, Deerfield, IL) and injected into the containers using a syringe to a final concentration of 10 μL/mL. This concentration was monitored by gas chromatography every 6 h and remained stable throughout the treatment. Control plants were incubated in sealed containers without ethylene injection. The containers were opened after 24 hours, leaf samples (300 mg) were collected and processed for total RNA isolation. We also measured ethylene emission in 90-day-old HP and WT plants by gas chromatography.

### Gene expression analyzes through quantitative Real Time PCR (qRT-PCR)

For the circadian experiment, plants were grown in a 12 hours light / 12 hours dark regime in MS20 medium at 24°C. Samples were harvested each six hours for 48 hours as follow: 12 PM – midpoint of the light period; 6 PM – start of the dark period; 12 AM – midpoint of the dark period; 6 AM – start of the light period. Each sample comprised a pool of four whole seedlings. All the other qRT-PCR experiments were done using plants grown under a 14-hour photoperiod at 24°C.

RNA was extracted from samples using a modified LiCl method [60] and treated with DNaseI (Ambion). cDNA was synthesized using SuperScript First-Strand Synthesis System for RT-PCR (Life Technologies) following the manufacturer’s instructions. The real time RT-PCR reactions were performed using the SYBR Green Real-Time PCR Master Mix according to manufacturer’s instructions. All experiments were based on three biological replicates with three technical replicates each.

## Supporting information

Hernandes-Lopes & Quintanilha et al figure additional figures

## LIST OF ABBREVIATIONS

cDNA: complementary DNA
CPM: Counts per million
DEGs: Differential expression genes
EFEs: Ethylene forming enzymes
FDR: False discovery rates
Gag: Group antigens
GO: Gene ontology
GSI: germination seed index
HP: Hair-pin lines
INT: Integrase
LTR: Long terminal repeats
MCE: Mean conditional entropy
MGT: Mean germination time
miRNA: micro RNA
mRNA: messenger RNA
ORFs: Open reading frames
PR: pathogenesis-related
qPCR: Quantitative polymerase chain reaction
RNAi: RNA interference
RPKM: Reads per kilo base per million mapped reads
RT: Reverse transcriptase
SGN: Sol genomics network
sRNA: small RNA
TE: Transposable Elements
TMM: Trimmed means of M values
WT: Wild type

## DECLARATIONS

## Ethics approval and consent of participate

Not applicable

## Consent for publication

Not applicable

## Availability of data and materials

Raw sequences generated in this study are deposited in GenBank (https://www.ncbi.nlm.nih.gov/genbank/) as GEO accession number GSE44027.

## Author’s contributions

Conceptualization MAVS, DQ and JHL; Data curation DQ, EMJ and JHL; Formal analysis DQ, EMJ, JHL, FL, JPK, MAVS; Funding acquisition MAVS; Methodology AAC, RBP, FL, DQ, EMJ, JHL, HMD, BK; Supervision MAVS; Roles/Writing - original draft DQ, JHL, EMJ; Writing - review & editing AAC, TBJ, JHL, MAVS.

## Acknowledgements

We thank Dr. Tatiana Corrêa for assistance with experiments, Dr. Myna Nakabashi for technical support with plant transformation, Dr. Anna Christina de Mattos Salim for technical assistance with transcriptome sequencing and Dr. Luciano Freschi, Dr. Hana Masuda, Dr. Nathália de Setta, Dr. Walter Colli, Dr. Françoise Simon-Plaz and Dr. Marie-Angele Grandbastien for discussions and comments on the manuscript.

## ADDITIONAL FILES

**Additional file 1: Figure S1**. Tnt1 retrotransposon structure and RNAi construct used to interfere with Tnt1 levels of transcripts. **Figure S2.** Expression of circadian clock genes and GO categorization of modulated genes in HP lines. **Figure S3.** Diagram of a gene regulatory network observed exclusively in HP plants. **Figure S4.** Phenotypes found in roots of HP lines. **Figure S5.** Germination performance of fresh and 6-years-old seeds.

