## Supplementary material for "Evidence-based gene expression modulation correlates with transposable element knock-down": Hernandes-Lopes & Quintanilha et al figure additional figures

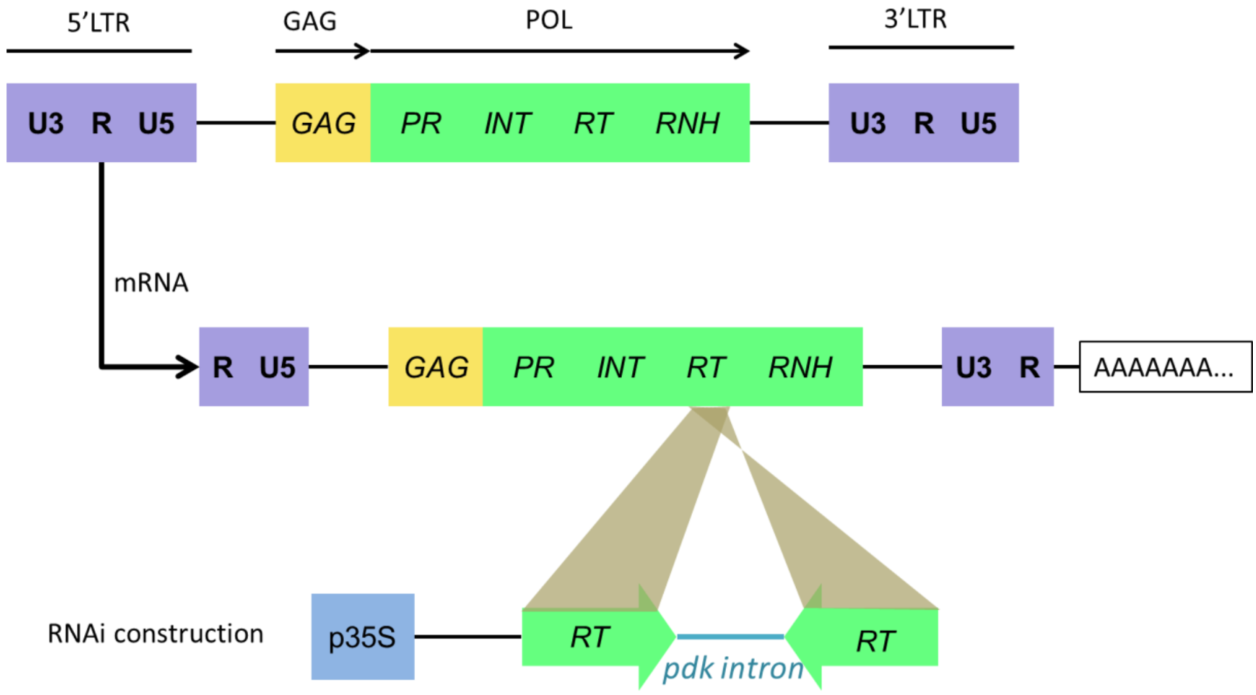


**Figure S1.** Tnt1 retrotransposon structure (top row), with 5´ and 3´ LTRs divided in U3 (unique 3’) R (repeated RNA) and U5 (unique 5’); and the protein domains contained in the element, capsid-like protein (GAG), protease (PR), integrase (INT), reverse transcriptase (RT) and RNase H (RNASEH). Middle row represents the Tnt1 transcript, which initiates at 5´ end of the R in the 5´ LTR and terminates at 3´ end of R in the 3´ LTR. Bottom row depicts the RNAi construct used to interfere with Tnt1 levels of transcripts. The construct consists of part of the RT domain in both sense and antisense directions separated by the *pdk* intron, which expression is driven by the p35S promoter.

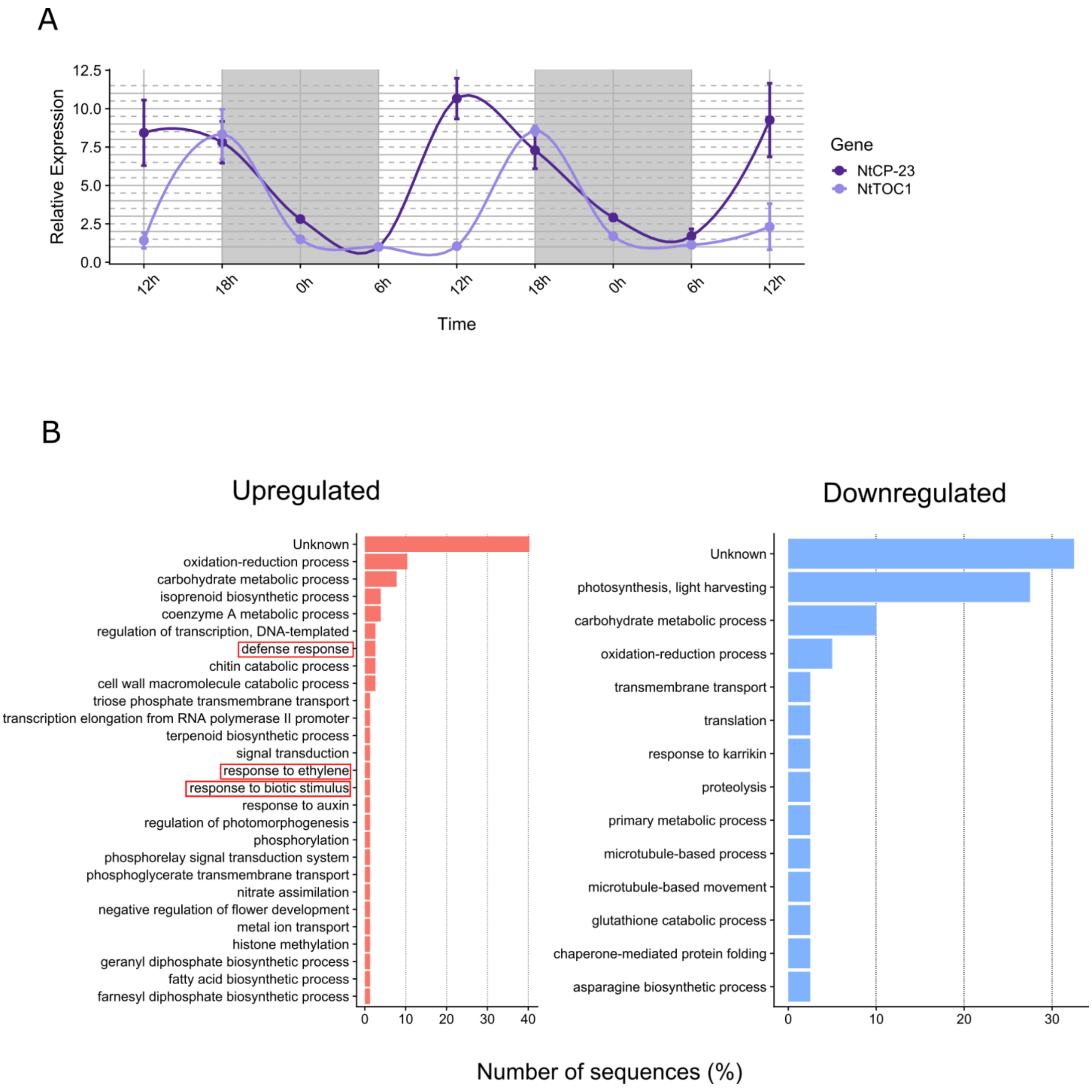


**Figure S2.** (A) Relative expression of circadian clock genes in 15-days-old WT plants throughout 48 hours period. Plants were grown under a 12 hours light / 12 hours dark regime. Gray areas indicate dark periods. The lowest expression value for each gene was set to one. (B) Gene ontology (GO) categorization of the 97 modulated genes found in HP leaves under a p-value of ≤ 0.01. Red boxes indicate stress-related processes.

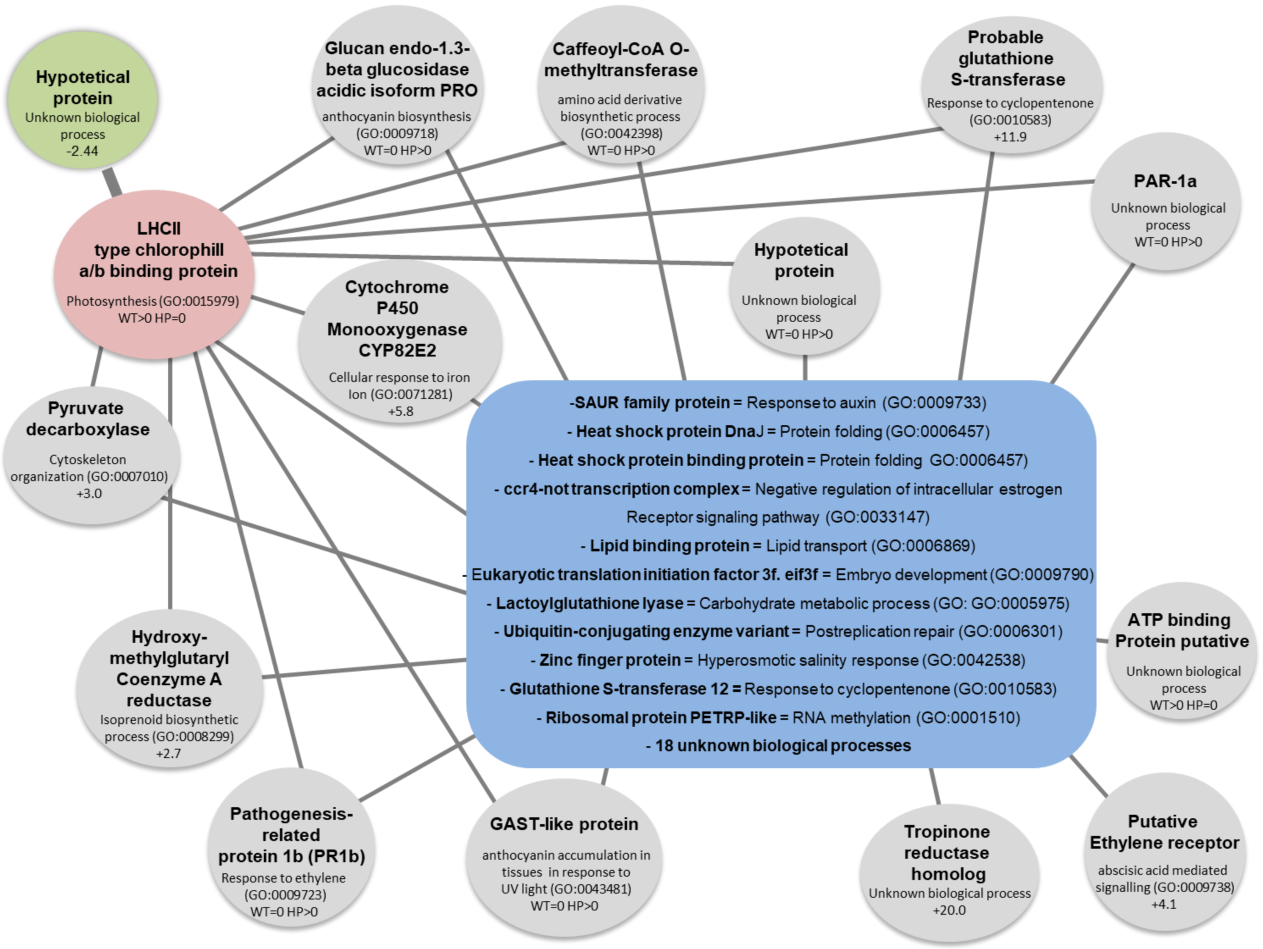


**Figure S3.** Diagram of a gene regulatory network observed exclusively in HP plants. The network is composed of forty-four genes of which we show fifteen. Each node (green. red and grey ellipses) denotes a gene. The blue box gathers a cluster of genes that connects to the red and grey nodes. From these eleven have a known biological function and eighteen which function is still unknown. The black lines that connect the genes represent a statistically supported link between changes in expression. Each node presents the gene annotation (bold), the major biological process to which it is related and the fold change. WT=0 HP>0 means induction of the gene in HP, with no detectable expression in WT. WT>0 HP=0 means repression of the gene in HP, with detectable expression in WT.

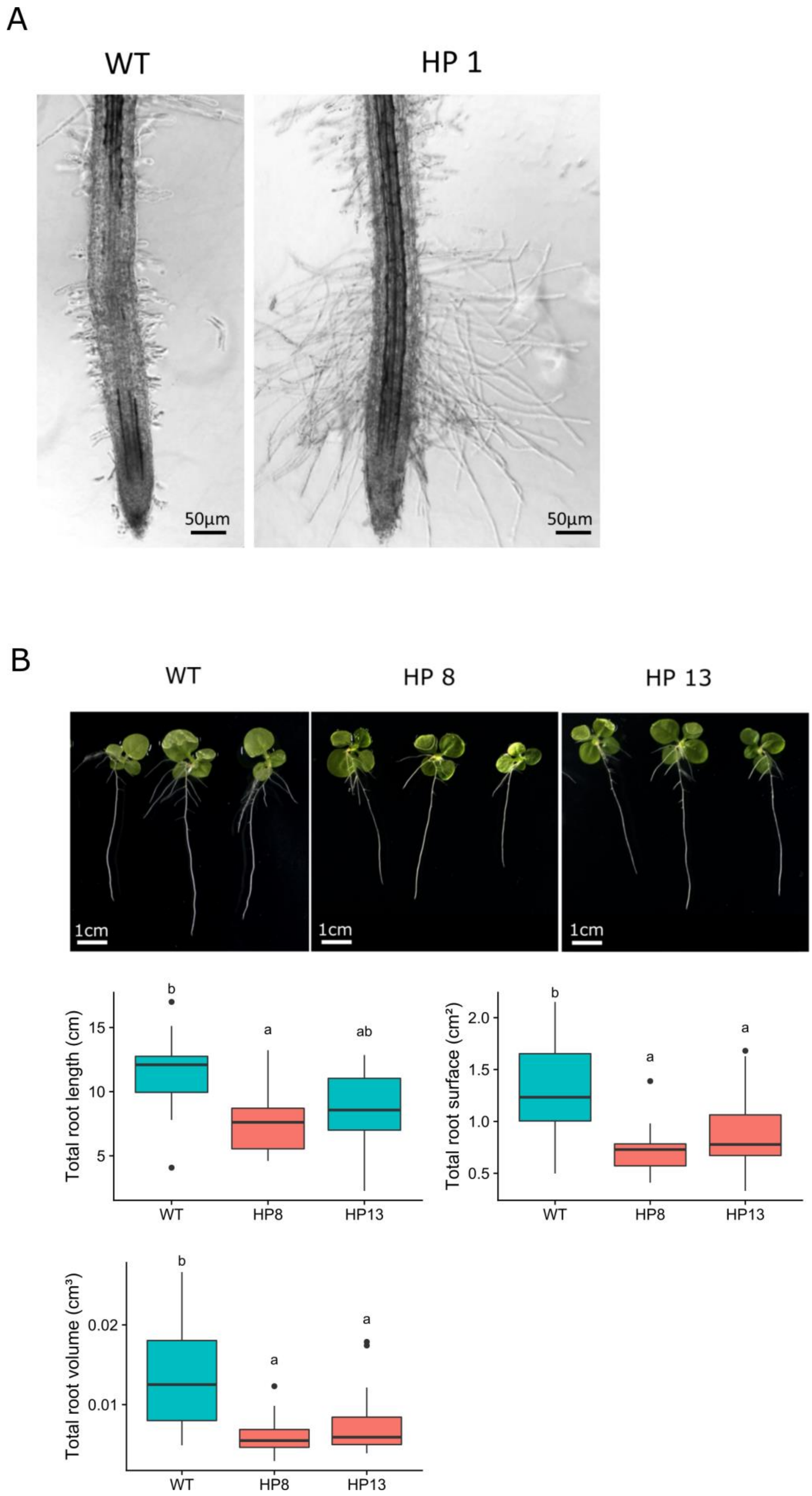


**Figure S4.** Phenotypes found in roots of HP lines. (A) Increased root hair growth in HP1 line. (B) Comparison between the root system of 15-days-old WT and HP lines. Letters indicate statistically significant differences between lineages.

### 6 years old seeds (T3)

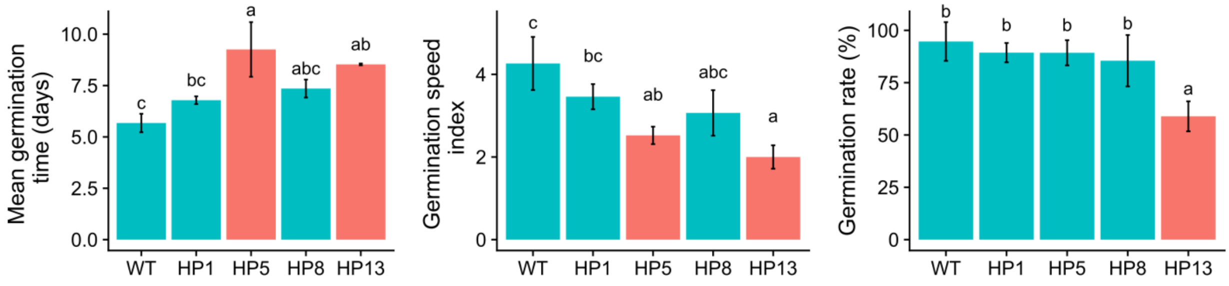


### Fresh seeds (T5)

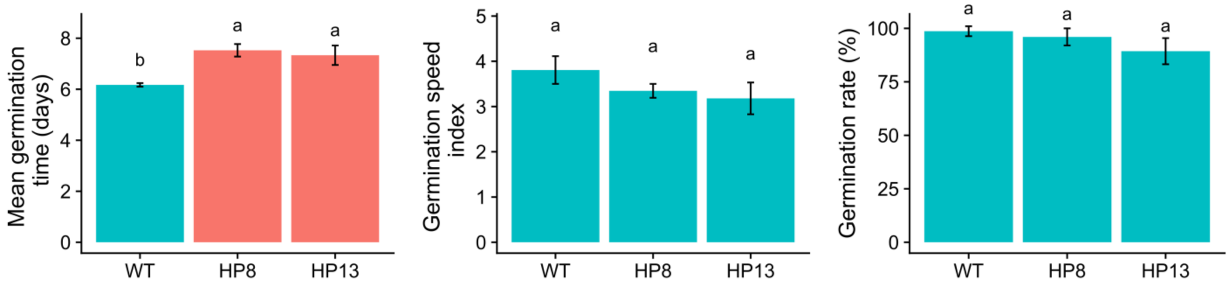


**Figure S5.** Germination performance of fresh and 6-years-old seeds. Mean germination time, germination speed index and germination rate of WT and HP lines. Letters indicate statistically significant differences between lineages. Red bars indicate samples significantly different from WT.
